# Evolutionary and functional diversification of cork oak NLRs reveals RNL expansion and dual roles in biotic and abiotic stress

**DOI:** 10.64898/2026.04.20.719699

**Authors:** L.M. Gonçalves, M.M. Oliveira, P.M. Barros

**Affiliations:** Instituto de Tecnologia Química e Biológica António Xavier, Av. da República, 2780-157 Oeiras, Portugal

**Keywords:** Quercus suber, NLR genes, Plant immunity, InterNLR, Drought, Biotic stress, Positive selection, Balancing selection

## Abstract

The Nucleotide-Binding Domain Leucine-Rich Repeat (NLR) gene family is a central component of plant immune systems, yet its diversity and evolutionary dynamics remain poorly characterized in long-lived tree species. Here, we performed a genome-wide analysis of the NLR gene family in *Quercus suber* (cork oak) using InterNLR, a new annotation tool, and explored their expression regulation in response to biotic and abiotic stresses. A total of 918 NLR and NLR-like genes were identified, encompassing both canonical and non-canonical members. Phylogenetic analyses based on the NB-ARC domain highlighted the distinct evolutionary trajectory of RNL proteins, which function as helper NLRs and show evidence of clade-specific gene duplication. Transcriptomic analyses revealed pronounced tissue-specific expression patterns, with RNLs exhibiting significantly elevated expression in xylem, suggesting a specialized role in this tissue. Under drought stress, seven NLR genes were differentially expressed and shared orthology with known abiotic stress–responsive genes. Notably, a CNL gene (LOC111996439) responded to both biotic and abiotic stresses, indicating a potential role as an integrative regulator of early defence responses, while an ADR1 orthologue (LOC112022539) suggests molecular crosstalk between stress signaling pathways. Population genetic analyses further revealed signatures of both positive and balancing selection acting on NLR genes. Together, these results provide new insights into the evolution, expression, and functional diversification of NLRs in cork oak. This work advances our understanding of immune gene architecture in an ecologically and economically important forest tree species.

## Introduction

Rising temperatures, altered precipitation patterns and the increasing frequency of extreme weather events are examples of environmental shifts that bring significant challenges to ecosystems worldwide and therefore implications for plant survival (Travis, 2003). Cork oak (*Quercus suber* L.), a sclerophyllous evergreen tree species from the Fagaceae family, is particularly affected, mostly due to its long lifespan (Pina-Martins et al., 2018). This species is mostly distributed in the Western Mediterranean region, where it plays an ecologically important role by maintaining the biodiversity, supporting soil stability and contributing to carbon sequestration (Hidalgo et al., 2008). Cork oak has also an important role by sustaining traditional agroforestry systems, such as *montado* and *dehesa*, which are crucial for local economies and cultural heritage (Besson et al., 2014).

Drought is one of the greatest challenges for the survival of cork oak stands since water stress events impose a significant threat to tree vitality (Cook et al., 2018). During drought adaptation, water uptake from the roots and transport capacity in the xylem is reduced and affects the overall hydraulic function of the tree (Chaves et al., 2003). This results in decreased tree growth, wilting, and even mortality of cork oak trees. Some studies have shown that drought conditions can alter the anatomy and physiology of xylem vessels – reducing vessel size and increasing embolism formation – therefore compromising water transport efficiency (Camilo-Alves et al., 2017; Besson et al., 2013). Drought can also reduce tree defences, increasing susceptibility to pathogens, as observed in other tree species (McDowell et al., 2008). In the case of cork oak, drought has been shown to enhance the vulnerability of the trees to pathogens induced by stress such as the fungus *Hypoxylon mediterraneum* (Vannini et al., 1996) and the oomycete *Phytophthora cinnamomi* (Moreira et al., 2006). Therefore, it is important to understand the interplay between drought and biotic stress for effective forest management and conservation strategies.

In response to biotic stimuli, plants rely on their innate immune system to defend themselves from pathogen infection. Many pathogens deliver effector proteins to the intracellular environment to manipulate host metabolism. In turn, host plants have developed a second level of surveillance and effector recognition, the effector-triggered immunity, which is mediated by intracellular immune receptor proteins named Nucleotide Binding and Oligomerization domain (NOD)-like receptors, also known as Nuclear-binding Leucine-rich-repeats (NLRs). These receptors are related to specific innate immunity receptors in animals.

After detecting specific effectors, NLRs trigger a hypersensitive response that may lead to cell death at the infection site, thus preventing the expansion of the pathogen (Wu et al., 2017).

NLRs are multi-domain proteins with a conserved modular architecture. Canonically, they comprise a variable N-terminal domain, which often functions as a signalling domain, a central nucleotide-binding (NB) domain that contains two ARC domains, and a C-terminal leucine-rich repeat (LRR) domain that plays a role in sensing ligands and potentially also in activity regulation (Jones et al., 2016). NLRs detect effector proteins secreted by pathogens through direct or indirect interactions. For example, they can directly bind effector proteins or indirectly interact with them through effector-targeted host proteins. An emerging model suggests that some “sensor” NLRs, responsible for detecting pathogen effectors, require “helper” NLRs to initiate immune signalling. Helper NLRs are more efficient at monitoring targets of effectors than the effectors themselves (Cesari, 2018).

Depending on the N-terminal domain, NLRs are classically classified as TNLs (with a Toll-interleukin receptor, or TIR domain), CNLs (with a Coiled coil) or RNLs (with a RPW8, Resistance to Powdery Mildew 8). However, in recent years, numerous studies have revealed that NLRs are more diverse than previously thought and variable in both their structure and their mode of action and there is evidence of truncated versions of NLRs playing a functional role in biotic stress response. For example, the TIR-only protein RBA1 recognizes a *Pseudomonas syringae* effector and regulates cell death in *Arabidopsis* (Nishimura et al., 2017). It is still unknown how truncated NLRs remain inactive in the absence of a pathogen, considering that they lack the canonical domains usually involved in the control of NLR activity - the NB and/or the LRR (Cesari, 2018).

While NLRs are primarily associated with biotic stress responses, emerging evidence suggests their involvement in abiotic stress tolerance (Ariga et al., 2017; Rizwan et al., 2023). Ghelder et al. (2019) explored the response of conifer *NLR* genes to drought stress. Their analysis revealed that different groups of *RNLs* exhibited varying responsiveness to drought conditions. Notably, certain *RNLs* demonstrated significant overexpression in response to drought. In other studies, some RNL proteins like Activated Disease Resistance 1 (ADR1) were required for the signaling of three different Arabidopsis NLRs recognizing effectors. Interestingly, constitutive or conditional enhanced expression of ADR1 conferred significant drought resistance in *Arabidopsis* (Chini et al., 2004).

The objective of this study was to identify and characterize the full repertoire of NLR genes in the cork oak genome, and investigate their potential roles in the response to environmental stress. Both canonical and non-canonical NLRs were identified, encompassing diverse domain architectures containing at least the NB-ARC domain. To infer their potential function we examined gene expression patterns across different cork oak tissues from plants exposed to drought conditions or infected with *Phythophthora cinnamomi*. Finally, using RNA-seq data, we explored the evolutionary dynamics of the NLR repertoire by analysing gene structure and the genetic diversity in two Iberian cork oak populations.

## Results

### Identification and characterization of NLR genes in cork oak and related species

Understanding the diversity of nucleotide-binding leucine-rich repeat receptors (NLRs) is essential to further explore their roles in plant immune response. For the identification of the cork oak *NLR* gene family we developed InterNLR, a tool which categorizes each NLR gene, as either CNL, TNL, or RNL, based on its domain structure and respective order. We identified a total of 374 NLRs in cork oak - 163 TNLs, 193 CNLs, and 18 RNLs. The ratio “number of RNLs” to “number of TNLs” stood at 0.1104. The same procedure was repeated for translated CDS sequences from *Quercus lobata* and *Quercus robur* to further explore the applicability of InterNLR. A total of 408 canonical NLRs were identified for *Q. lobata* - 199 TNLs, 197 CNLs, and 12 RNLs - and 358 canonical NLRs for *Q. robur* - 165 TNLs, 177 CNLs and 16 RNLs. The ratio “number of RNLs” to “number of TNLs” was 0.0603 and 0.0969 for *Q. lobata* and *Q. robur*, respectively.

We conducted a maximum likelihood phylogenetic analysis based on the NB-ARC domain of all cork oak canonical NLRs (Figure 1). We then expanded this analysis to include NLRs from all three *Quercus* species (Supplementary Figure 1). Specifically looking at cork oak NLRs, the phylogenetic study revealed a clear and distinct separation of CNL, TNL, and RNL clades. However, we did observe two NLRs that showed a discordant placement according to their predicted family, namely LOC111989151 (CNL in the RNL cluster) and LOC112034384 (TNL in the CNL cluster). This result was consistent in the phylogenetic analysis of NLRs from the three *Quercus* species, using the conserved NB-ARC domain (Supplementary Figure 1). Furthermore, we detected evidence of gene duplication within cork oak, particularly evident in the RNL group (Supplementary Figure 1). Genes such as LOC112040545 and LOC112040547 exemplify this phenomenon.

**Figure 1:**
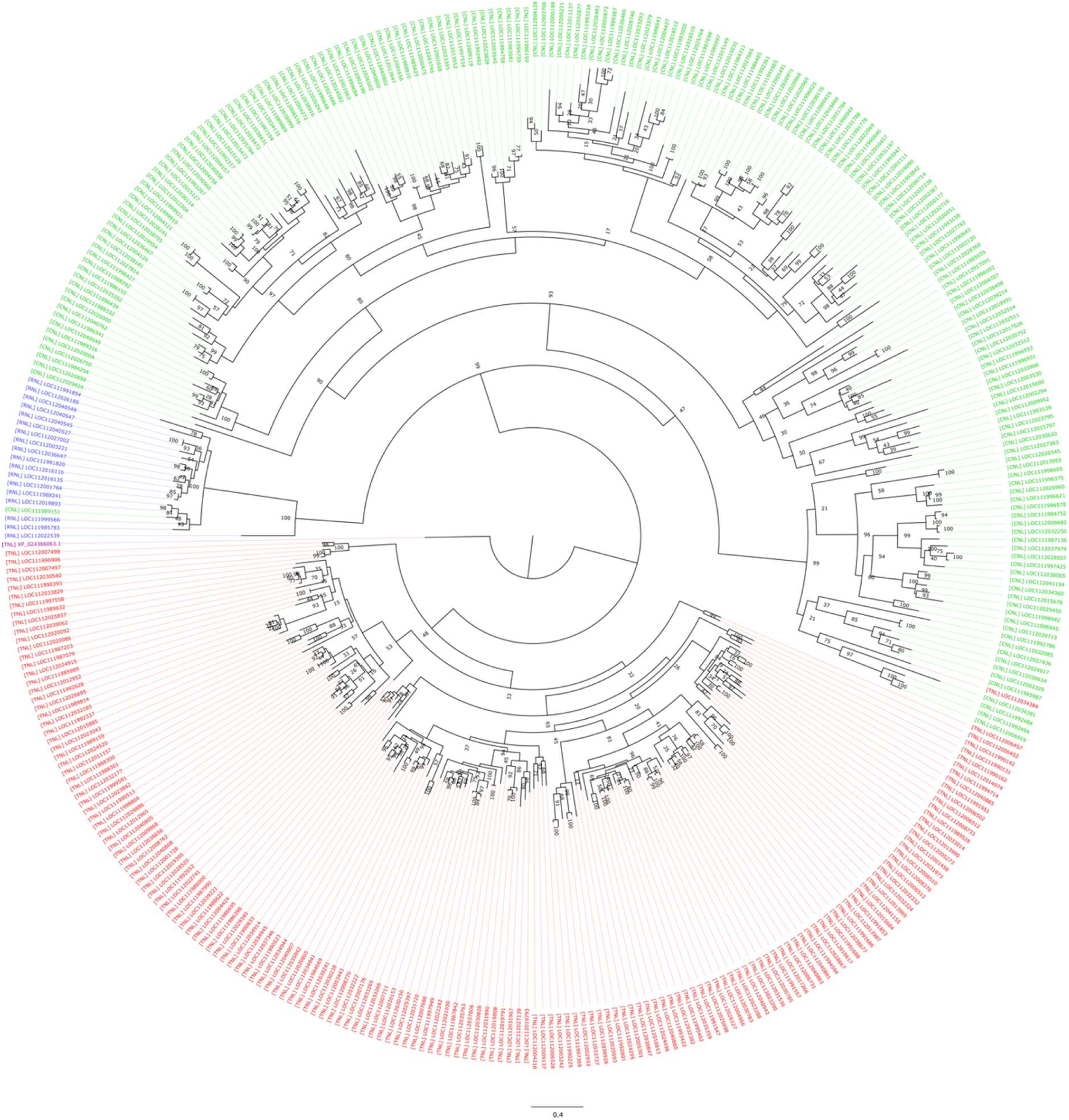
Maximum Likelihood Phylogeny of cork oak NLRs based on NB-ARC domain. Colors indicate NLR sub-groups: Red – TNLs; Green – CNLs; Blue – RNLs. The NB-ARC domain of *Physcomitrium patens* PRG gene was used as outgroup (purple). It is possible to distinct the polarization of each of CNL, TNL and RNL clades. The phylogenetic tree scale represents substitutions per site. Branch support values were generated from bootstrap analysis (1000 replications).

To elucidate the gene duplication patterns within cork oak NLRs as compared to those of *Q. lobata* and *Q. robur*, we employed Orthofinder (Emms et al., 2019) to conduct separate analyses for TNLs, CNLs, and RNLs (Supplementary Tables 1, 2, and 3, respectively). When analyzing TNLs, we identified 22 orthogroups in which gene duplications were detected. Regarding CNLs and RNLs, we pinpointed 16 and one orthogroups, respectively (Supplementary Table 4).

In addition to the traditional NLRs previously identified, our approach also uncovered NLRs with every possible domain combination and with at least NB-ARC, broadening the spectrum of NLR gene diversity. This raised the total number of NLRs to 918 in cork oak. Alongside the canonical NLRs identified, we found 230 NLs (lacking an N-terminal domain), 143 Ns (lacking both N-terminal and C-terminal domains), 10 TCNLs (with TIR and CC domains), 1 TRNL (with TNL and RPW8 domains), 60 TNs (only with TIR and NB-ARC domains), 93 CNs (with CC and NB-ARC domains) and 7 RNs (RPW8 and NB-ARC domains only) (Supplementary Figure 2, panel a). Phylogenetic analysis including all cork oak NLRs (canonical and non-canonical) again grouped the genes in three main clades, identified as TNL-like, CNL-like, and RNL-like (Supplementary Figure 2, panel b). Ratios “number of TNLs” to “TNL-like genes”, “CNLs” to “CNL-like genes” and “RNLs” to “RNL-like genes” were, respectively, 0.5, 0.350 and 0.45. A total of 780 and 867 NB-ARC-containing canonical and non-canonical NLRs were identified with InterNLR in *Quercus robur* and *Quercus lobata* genomes, respectively, without considering splicing variants.

### Drought-responsive NLR expression profiles in cork oak stems

To identify candidate NLR genes with a potential role in abiotic stress response in cork oak stems, we used a publicly available RNA-seq dataset obtained from outer bark (phellem), phloem and cortex (inner bark) and xylem from 1.5 year-old plants (Barros et al. 2024). This dataset included plants grown under well-watered (WW) and water-deficit (WD, controlled dehydration for 2 months followed by 4 months at 10% relative water content) conditions, which allowed us to assess transcriptional dynamics of NLRs in response to drought.

We first compared the average expression levels of the three canonical NLR gene classes with active expression in stem tissues. While statistically significant differences were not detected in mean expression among the NLR classes in both phellem and inner bark tissues, in xylem we detected statistically significant increased average expression of RNLs when compared to all other NLR classes except TRNL (Figure 2). Additionally, there was a higher proportion of transcribed canonical NLRs compared to non-canonical NLRs (Supplementary Table 5). The top five NLRs with the highest expression values are indicated in Table 1, with LOC112003710 (RPP13) showing the highest representation in phellem, while LOC112022539 (ADR1-L1) showed high expression levels in inner bark and xylem. To further assess tissue specificity of NLR genes we analyzed the diversity of differentially expressed NLRs between the three tissue layers, independently of the growth conditions. From the hierarchical clustering based on mean expression per biological replicates (Figure 3), we found: 39 canonical and non-canonical NLRs enriched in phellem (including 7 TNLs, 12 CNLs and no RNLs), with some of these also represented in inner bark; 26 canonical and non-canonical NLRs enriched in inner bark (including 5 TNLs, 5 CNLs and no RNLs); and 52 canonical and non-canonical NLRs more expressed in xylem (including 15 TNLs, 15 CNLs and 1 RNLs).

**Figure 2:**
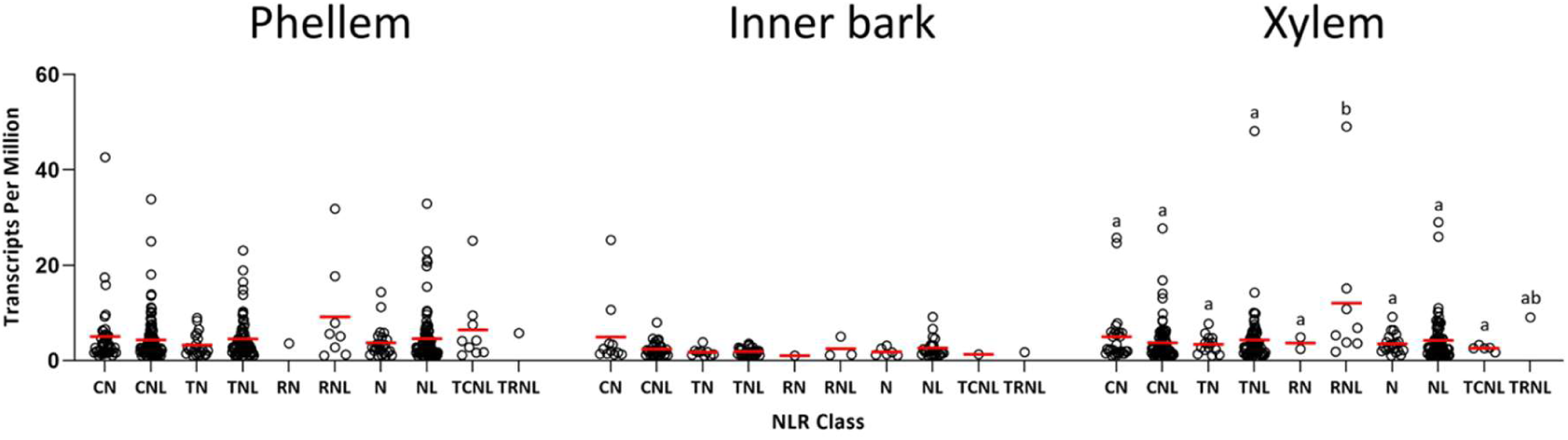
Dot plot representing the gene expression values (TPM) grouped according to NLR class and stem tissue. Red lines represent the mean TPM value. Different lowercase letters indicate a statistically significant difference (p < 0.05) as determined by one-way ANOVA followed by a Tukey post-hoc test. Groups that share at least one letter are not significantly different.

**Figure 3:**
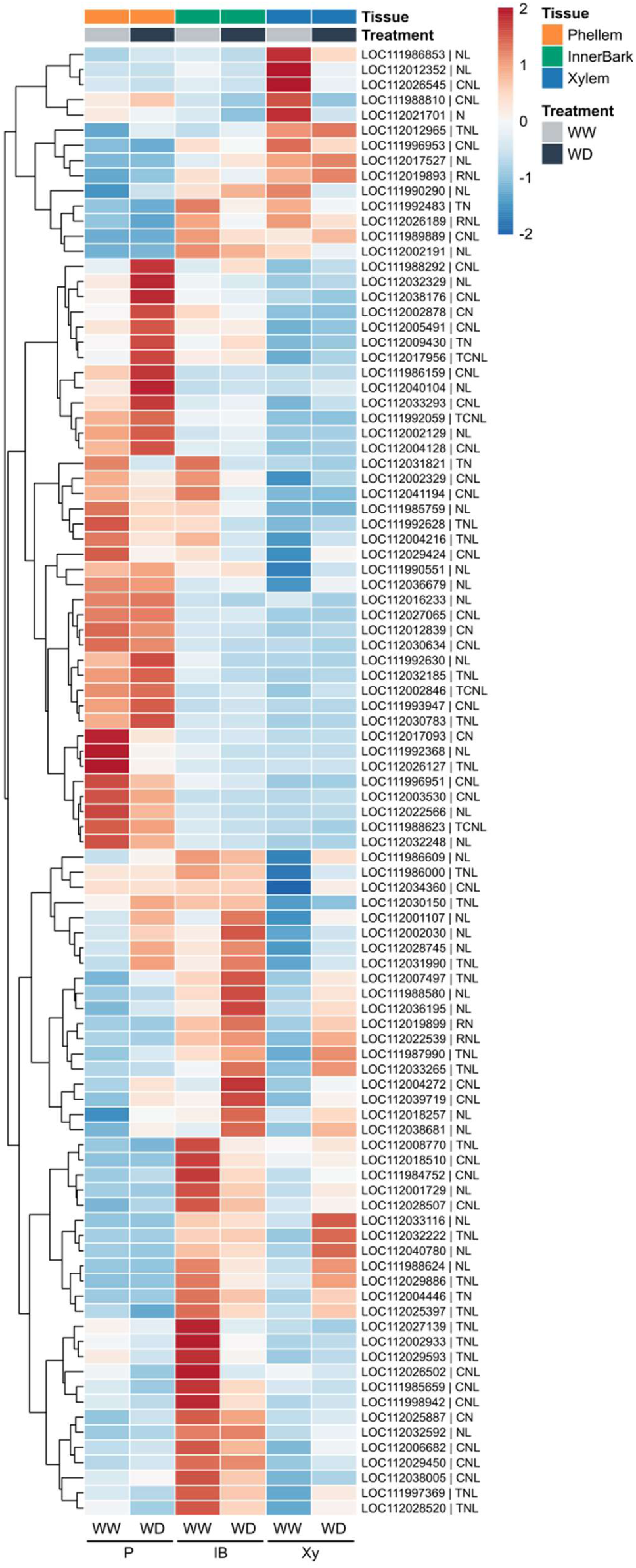
Gene expression heatmap for NLR genes differentially expressed between stem tissues from 1.5 year-old plants (Likelihood Ratio Test, adjusted p-value < 0.001), independently of growth condition. Gene expression is represented by Z-scores of the TPM values to enable comparison of relative expression across genes and samples. WW, Well Watered; WD, Water Deficit; P, Phellem; IB, Inner Bark; Xy, Xylem.

**Table 1:** Top five *NLRs* with higher mean expression in phellem, inner bark and xylem collected from 1.5 year-old plants. Expression values are represented in Transcripts Per Million. The corresponding *Arabidopsis* orthologue gene obtained using Blastp is also indicated.

| Gene name | NLR class | Mean Expression* |  |  | *Arabidopsis* orthologue | E-value |
| --- | --- | --- | --- | --- | --- | --- |
|  |  | Phellem | Inner Bark | Xylem |  |  |
| LOC112003710 | CN | (1) 42.62 | 31.91 | 25.75 | RPP13 | 1.14E-33 |
| LOC112033293 | CNL | (2) 33.82 | 19.65 | 13.05 | AT3G14470 | 1.81E-84 |
| LOC112022566 | NL | (3) 32.92 | 3.23 | 1.06 | AT5G17680 | 3.17E-113 |
| LOC112022539 | RNL | (4) 31.83 | (1) 73.39 | (1) 49.05 | ADR1-L1 | 0 |
| LOC112002846 | TCNL | (5) 25.11 | 5.24 | 2.53 | AT4G11170 | 1.07E-130 |
| LOC112036463 | NL | 19.81 | (2) 41.32 | (3) 29.01 | AT3G14470 | 1.22E-114 |
| LOC112036467 | CNL | 18.01 | (3) 40.29 | (4) 27.72 | AT3G14470 | 1.53E-133 |
| LOC111988332 | CNL | 10.01 | (4) 36.47 | 16.80 | AT3G14470 | 1.51E-127 |
| LOC112029450 | CNL | 7.58 | (5) 32.22 | 5.97 | AT4G27190 | 1.52E-71 |
| LOC112012965 | TNL | 18.88 | 24.21 | (2) 48.11 | AT5G17680 | 1.36E-154 |
| LOC112001962 | NL | 22.90 | 28.10 | (5) 25.94 | AT1G50180 | 3.03E-170 |

\*Numbers in parentheses indicate the ranking order for the top five for each tissue.

To explore the relationship between NLR expression and adaptation to drought, we looked for NLR genes in the list of differentially expressed genes found in WD vs. WW comparison per tissue (Barros et al. 2024). We identified 36 differentially expressed (DE) NLRs in the phellem (Supplementary Figure 2, panel d1), 29 in the inner bark (Supplementary Figure 2, panel d2), and 97 in the xylem (Supplementary Figure 2, panel d3). Among these DE NLRs, 9 genes found in both phellem and inner bark, 2 genes in both phellem and xylem, and one differentially expressed in both inner bark and xylem (Figure 4a). To gain deeper insights, we selected the differentially expressed NLRs for further analysis and TPMs were calculated for each tissue and condition (Figure 4b). A total of 25, 6 and 71 canonical and non-canonical NLRs were up-regulated on drought stress, respectively, in phellem, inner bark and xylem tissues. A total of 10, 23 and 25 were down-regulated, respectively, in the same three tissues.

**Figure 4:**
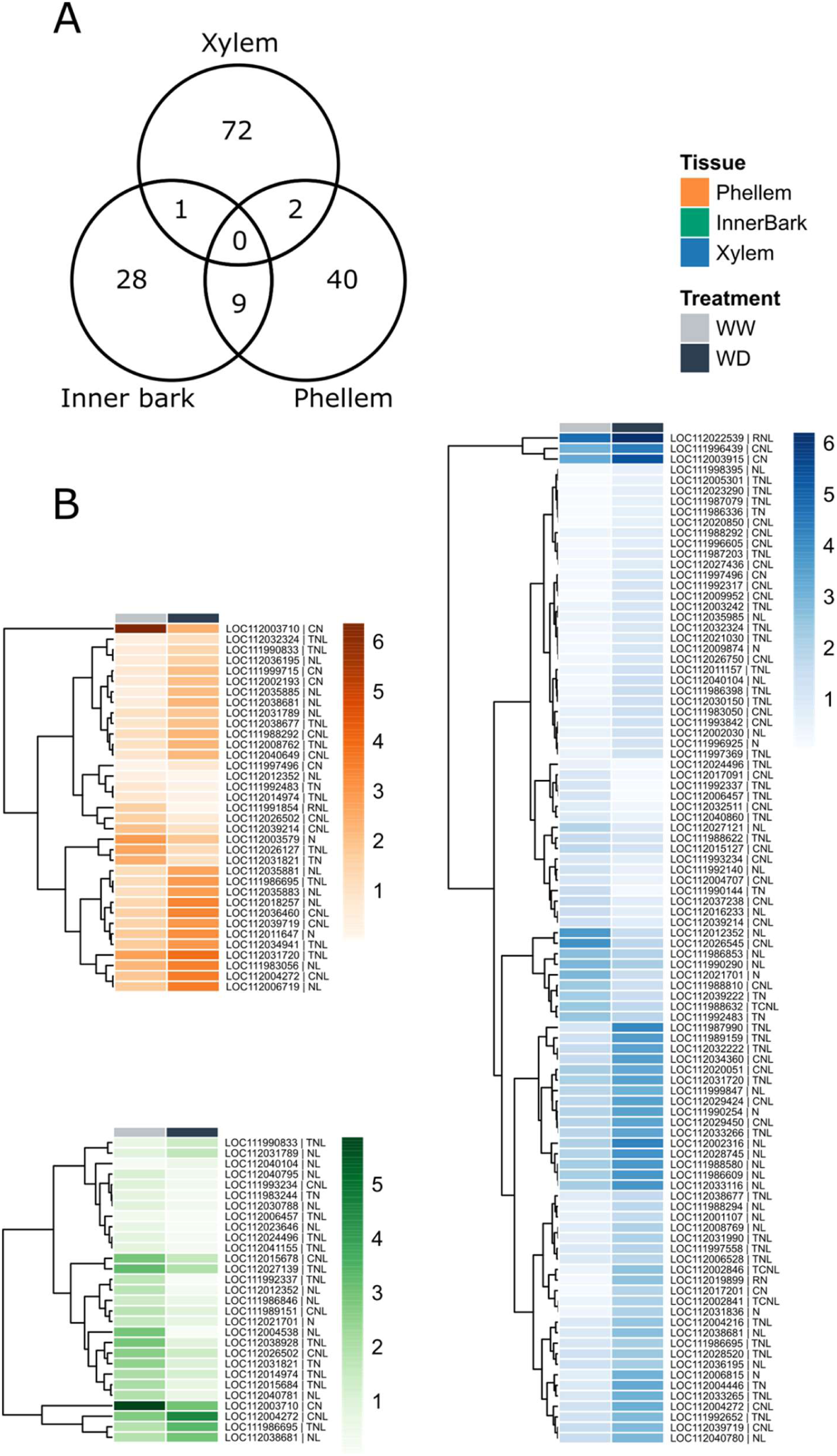
Differently expressed canonical and non-canonical NLRs identified comparing WD vs. WW conditions in phellem, inner bark and xylem from cork oak 1.5-year-old stems. A: Venn diagram of the intersection of the differentially expressed NLRs in phellem, inner bark, and xylem tissues. B: Gene expression heatmaps for DE NLRs identified for phellem, inner bark and xylem; WW – Well watered, WD – Water Deficit; Gene expression is represented by log_2_(TPM+1) values.

To further explore the list of DE NLRs we analyzed their homology with NLRs from other plant species which have been reported as drought responsive. A total of 27 cork oak NLRs (from canonical and non-canonical groups) showed high homology to NLRs from Arabidopsis (Ariga et al. 2017, Chini et al., 2004, Yang et al., 2021), *Arachis hypogea* (Rizwan et al. 2023), *Polygonatum kingianum* (Qian et al., 2021), *Oryza sativa* (Yang et al., 2022), *Nicotiana benthamiana* (Li et al., 2017) and *Picea glauca* (Ghelder et al., 2019), based on both best Blastp hit (with reciprocal best hit) and Orthofinder (Table 2; Supplementary Table 6). From these, 7 were differently expressed genes in at least one of the three stem tissues (Supplementary Figure 2, panel c; Table 3). These were LOC112031821 (TN), LOC112006457 (TNL), LOC112021701 (N), LOC111996925 (N), LOC111996439 (CNL), RPM1 (LOC112040104, NL) and ADR1-like (LOC112022539, RNL).

**Table 2:** NLRs orthologues to genes with proved relation to abiotic stress in other plant species.

| Gene name* | NLR classes | Mean expression (TPM) |  | Arabidopsis orthologue (Best Blastp hit) |  |
| --- | --- | --- | --- | --- | --- |
|  |  | In Drought transcriptomic assay | In *P. cinnamomi* transcriptomic assay | Gene name | E-value |
| LOC111990393 | TNL | 0.254 | 0.412 | AT1G27170 | 0 |
| LOC111987799 | TN | 0.882 | 0.000 | AT5G17680 | 1.11E-116 |
| LOC112035764 | N | 0.050 | 0.000 | AT5G17680 | 1.63E-70 |
| LOC112035762 | N | 0.256 | 0.000 | AT5G17680 | 1.37E-71 |
| **LOC112031821** | TN | 1.925 | 0.485 | AT5G17680 | 5.39E-135 |
| LOC112034944 | TNL | 0.850 | 0.000 | AT5G17680 | 9.60E-169 |
| LOC112035366 | NL | 0.173 | 0.000 | AT4G12010 | 1.71E-129 |
| LOC111995402 | TNL | 0.585 | 1.666 | AT5G17680 | 0 |
| **LOC112006457** | TNL | 0.276 | 1.416 | AT5G36930 | 0 |
| LOC112006452 | TNL | 6.185 | 1.122 | AT5G36930 | 0 |
| LOC111996235 | N | 0.168 | 0.000 | AT5G36930 | 8.09E-81 |
| LOC111996063 | N | 1.058 | 0.000 | AT5G36930 | 3.93E-71 |
| LOC111988305 | TN | 4.848 | 0.000 | AT5G17680 | 1.91E-99 |
| **LOC112021701** | N | 2.665 | 9.388 | CW9 | 3.02E-09 |
| LOC111992775 | N | 0.951 | 0.000 | AT5G43730 | 1.68E-55 |
| LOC111992786 | CNL | 3.188 | 0.927 | RPS5 | 2.04E-141 |
| LOC112030634 | CNL | 1.972 | 0.632 | AT4G27220 | 4.84E-94 |
| **LOC111996925** | N | 1.869 | 5.285 | AT4G19060 | 4.08E-51 |
| LOC111993947 | CNL | 2.091 | 1.161 | AT3G14470 | 0 |
| LOC112040068 | N | 1.131 | 0.786 | AT3G14470 | 8.87E-87 |
| LOC111988289 | NL | 0.633 | 0.346 | AT3G14460 | 1.08E-61 |
| LOC1112035352 | CNL | 3.186 | 0.372 | AT3G14470 | 3.64E-133 |
| **LOC111996439** | CNL | 15.816 | 2.748 | AT3G14470 | 1.05E-136 |
| LOC111990551 | NL | 16.099 | 14.267 | AT1G50180 | 0 |
| LOC111987192 | NL | 0.014 | 0.000 | RPM1 | 2.15E-142 |
| **LOC112040104** | NL | 3.116 | 1.213 | RPM1 | 9.22E-149 |
| **LOC112022539** | RNL | 51.428 | 45.665 | ADR1-L1 | 0 |

\*Genes in bold are also differently expressed in drought stress conditions.

**Table 3:**
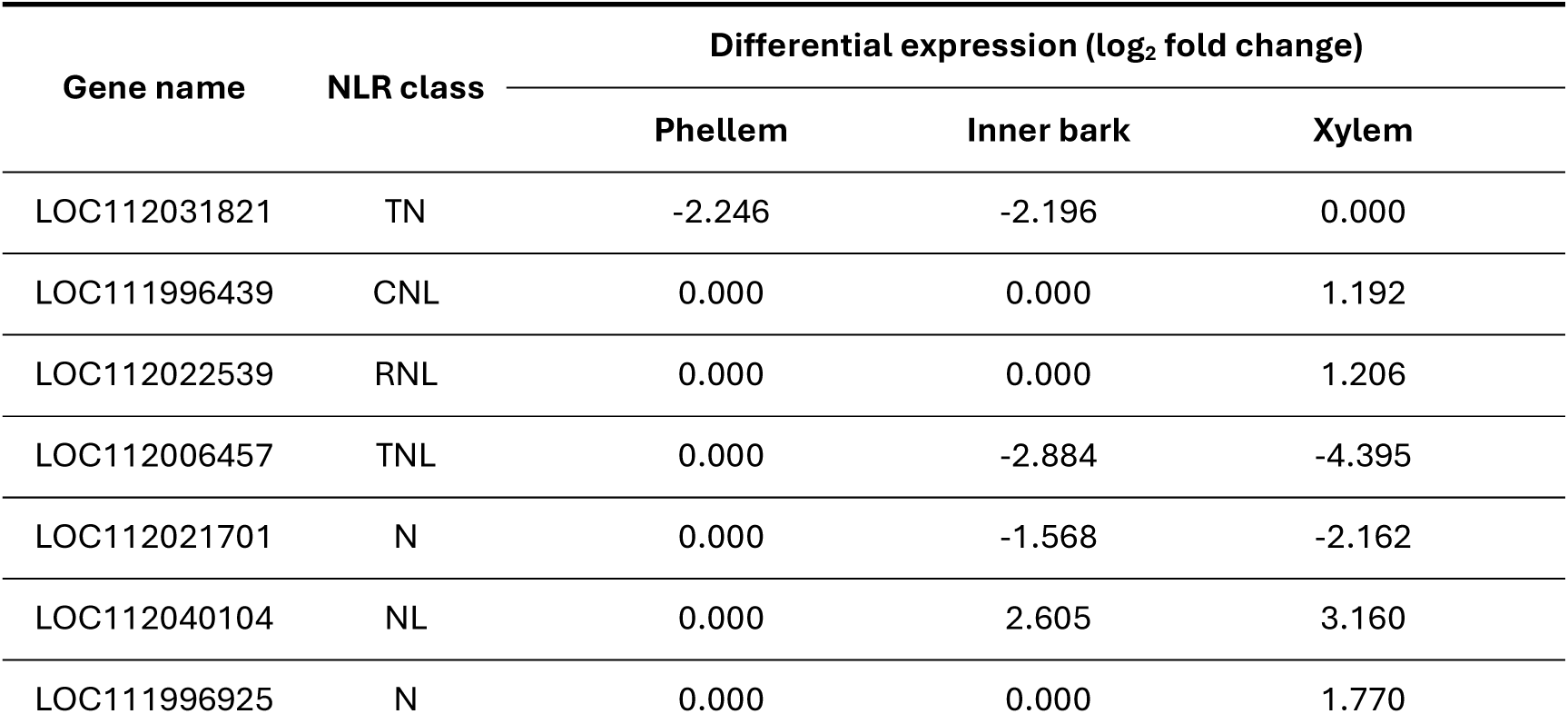
NLRs orthologues to reference ones with proved relation to abiotic stress and differentially expressed in drought stress transcriptomic assay.

### Expression analysis of NLRs in cork oak roots infected with *Phythophthora cinnamomi*

In order to gain insights into the potential functions of the identified cork oak NLRs, we made use of a publicly available transcriptomic dataset, retrieved from CorkOakDB (Arias-Baldrich et al. 2020), consisting of RNA-seq data of roots on different developmental stages - roots from germinated acorns (days old) and thin white roots (2 months old) - both infected and non-infected with *Phythophthora cinnamomi* zoospores.

We identified 168 genes displaying a minimum Transcript Per Million (TPM) of 1 across at least one tissue/condition (Figure 5). Additionally, we computed the average TPM of the transcriptomic assay, unveiling that LOC112022539 (ADR-L1), exhibited the highest value among them (Table 4). Included among the top 10 highest average TPM values were genes like LOC111992494, LOC112003710 and LOC112017093, which are orthologous to *Arabidopsis* RPS2, RPP13 and RPM1, respectively. From these, LOC112017093 and LOC111992494 are among the top five highest expression difference between infected and non-infected with *P. cinnamomi* zoospores in thin white roots (Table 5). The ratio is especially high for LOC112017093 case (20.085), suggesting that this gene is up-regulated in young roots when infected with the pathogen.

**Figure 5:**
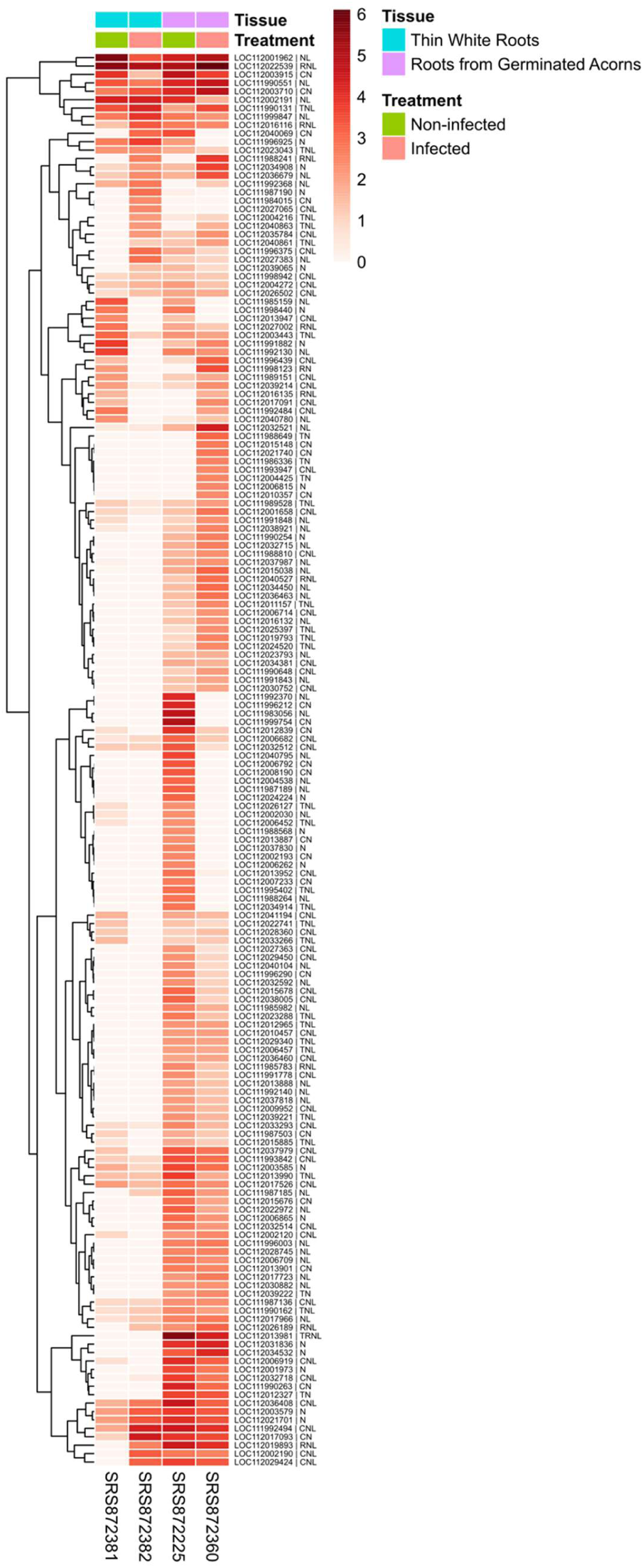
Gene expression heatmap for cork oak genes with strong expression in cork oak roots (TPM > 1) in at least one development stage and in response to *P. cinnamomi* infection. The heatmap scale represents log_2_(TPM+1) values.

**Table 4:**
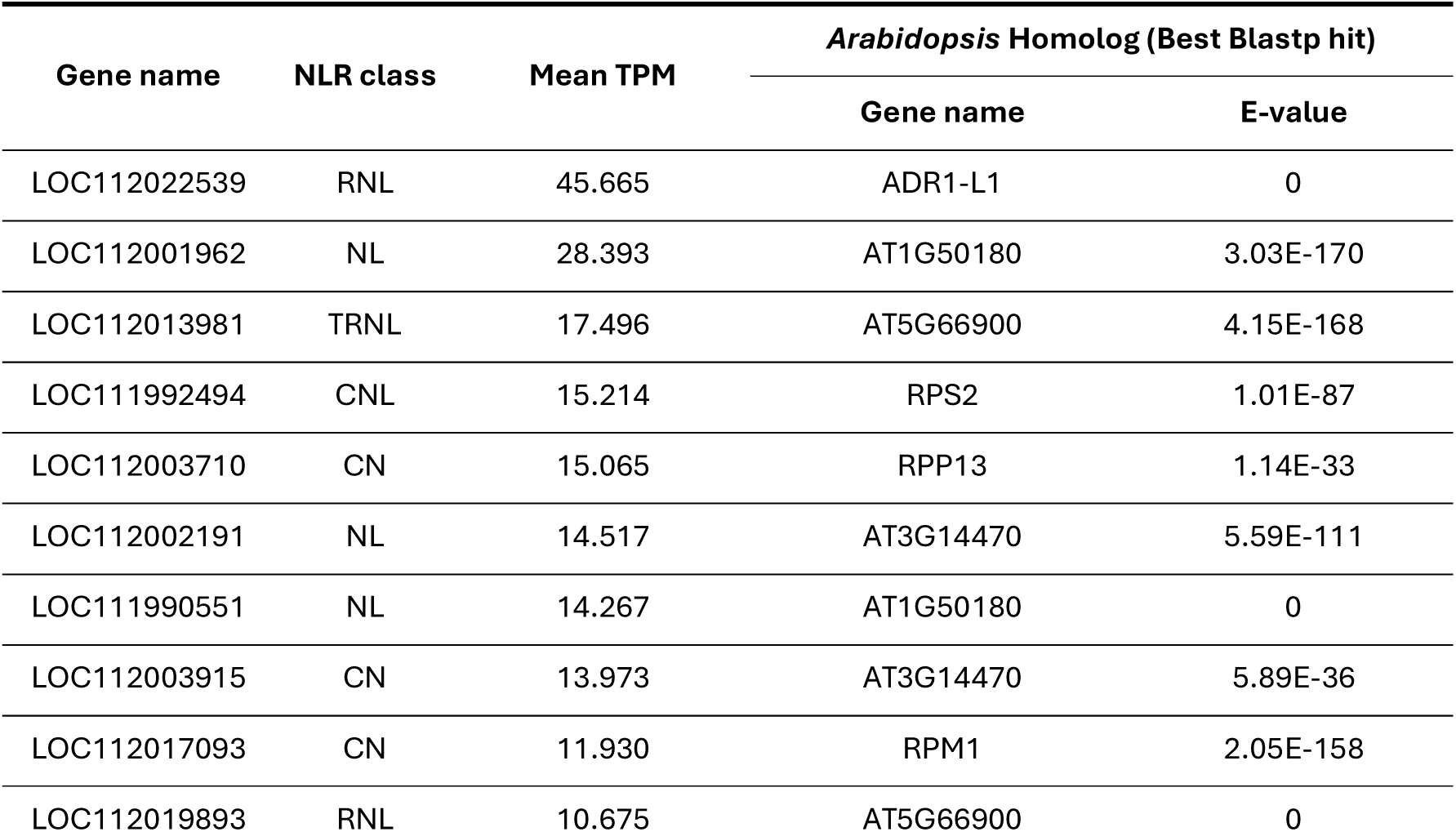
Top 10 NLRs with higher mean TPM values in cork oak roots infected with *Phytophtora cinnamomi* and control (non-infected) conditions.

**Table 5:**
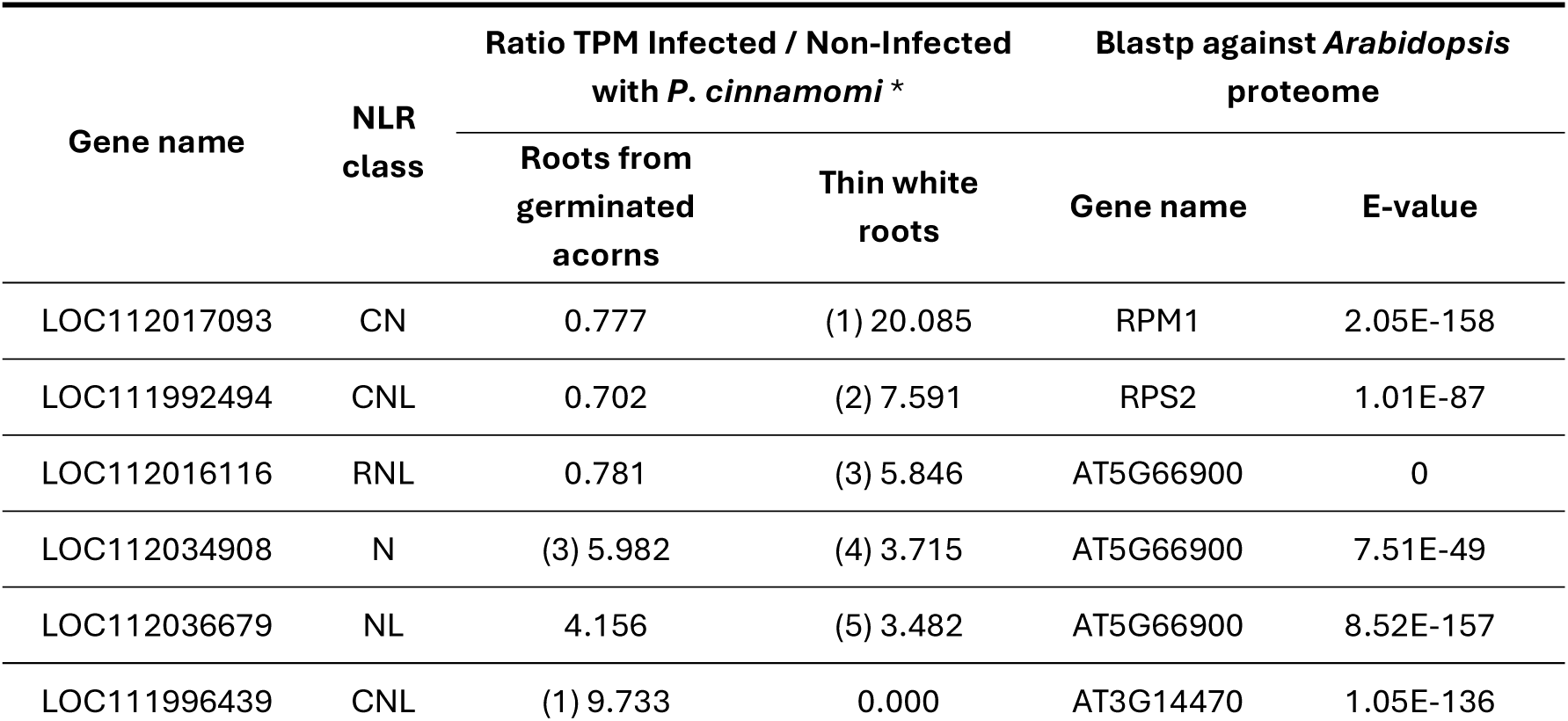
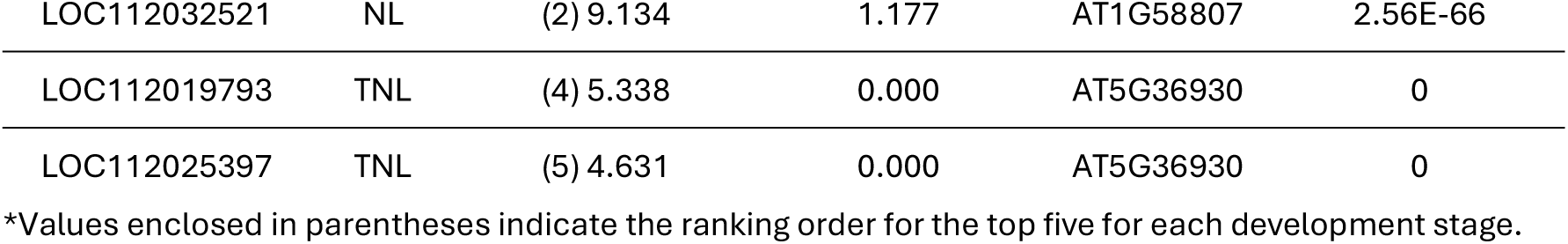
Cork oak NLR genes with the top five highest expression ratios of (based on TPM) in roots infected *versus* non-infected with *P. cinnamomi*.

### Population diversity analysis of NLRs

Addressing how NLRs behave in terms of genetic diversity is a step forward towards the understanding about their evolutionary dynamics. To evaluate genetic diversity of NLRs occurring in cork oak trees growing in the field, we used two independent RNASeq datasets obtained for individuals trees – 9 from Portugal (Population 1, Bioproject PRJEB33874, Lopes et al., 2019) and 11 from Spain (Population 2, Bioproject PRJNA650215, Fernández-Piñán et al., 2021). After read alignment against the reference genome, population diversity analysis was conducted using the information obtained for 564 NLRs (218 canonical and 346 non-canonical) displaying mean raw counts ≥10 reads across all the individuals. The final SNP filtering on the variant call file consisted on a depth of coverage ≥ 10 and a quality threshold ≥ 20. The weighted means for observed heterozygosity (H_O_), expected heterozygosity under Hardy-Weinberg Equilibrium (HE), nucleotide diversity (π) and Tajima’s D statistic were calculated for canonical and non-canonical NLRs for Portuguese (population 1) and Spanish (population 2) populations and having both populations into consideration (Table 6). Observed heterozygosity was lower than expected heterozygosity, and nucleotide diversity was low (order of magnitude of 10^-3^), for all cases. Tajima’s D values were positive, indicating a mean nucleotide diversity greater than the weight of the number of polymorphic sites (Watterson’s ϑ).

**Table 6:** Weighted mean of genetic diversity statistics across two Iberian cork oak populations, inferred based on commonly expressed genes from transcriptomic profiles of individual trees.

| Statistics<br>(weighted mean) | Population |  |  |
| --- | --- | --- | --- |
|  | Pop1 (PT) | Pop2 (ES) | Total (PT + ES) |
| $H_o$ | 0.2737 | 0.2974 | 0.2368 |
| $H_e$ | 0.3788 | 0.3740 | 0.3115 |
| $\pi$ | 0.0029 | 0.0016 | 0.0019 |
| Tajima's D | 1.0253 | 1.0418 | 1.0875 |

Next, we identified the top 15 (expressed) NLR genes showing the highest and lowest Tajima’s D scores. The top genes with highest Tajima’s D ranged between 1.587 and 2.623 and included mostly NLRs containing CC domain (9 out of 15). The 15 NLRs with lowest Tajima’s D scores, ranging between −0.476 and −1.049 included canonical and non-canonical NLRs from different groups. The 15 genes with the highest Tajima’s D showed greater absolute values than the 15 with the lowest (Table 7). Also, we identified NLRs with positive and negative Tajima’s D values, indicating distinct evolutionary trajectories. Furthermore, both groups of genes with top 15 highest and lowest Tajima’s D values include genes that are differentially expressed in response to drought - two being up-regulated in the top 15 with highest values, and another four being two down- and two up-regulated in the top 15 with lowest values.

**Table 7:** Top 15 NLRs with highest and lowest Tajima’s D values. Blue entries with light grey shading represent up-regulated genes in drought stress. Red entries with dark grey shading represent down-regulated genes.

| LOC ID | NLR type | Tajima's D |
| --- | --- | --- |
| LOC111990646 | CNL | 2.623 |
| LOC112027002 | RNL | 2.338 |
| LOC111990643 | CNL | 2.179 |
| LOC111992786 | CNL | 1.914 |
| LOC112031742 | N | 1.907 |
| LOC112017196 | N | 1.904 |
| **LOC112003221** | **RNL** | **1.867** |
| LOC112011137 | CNL | 1.849 |
| LOC112040625 | CN | 1.734 |
| LOC111986853 | NL | 1.709 |
| LOC111986155 | CNL | 1.677 |
| **LOC112018510** | **CNL** | **1.672** |
| LOC112003746 | CNL | 1.611 |
| LOC112017093 | CN | 1.606 |
| LOC112004425 | TN | 1.587 |
| LOC111993966 | NL | -0.476 |
| LOC112004336 | NL | -0.476 |
| LOC112005273 | TNL | -0.488 |
| LOC112033266 | TNL | -0.531 |
| **LOC112040007** | **TNL** | **-0.588** |
| LOC111993234 | CNL | -0.601 |
| LOC112004216 | TNL | -0.605 |
| LOC112017526 | CNL | -0.652 |
| LOC112026189 | RNL | -0.690 |
| LOC112040545 | RNL | -0.698 |
| **LOC112006502** | **TNL** | **-0.757** |
| LOC112004617 | NL | -0.960 |
| LOC112008190 | CN | -0.967 |
| **LOC112020850** | **CNL** | **-1.007** |
| **LOC112002030** | **NL** | **-1.049** |

Principal Component Analysis (PCA) based on NLR data explained 9.86% of the variance in PC1 (Figure 6). This analysis revealed a clear separation of Population 2 accessions from Population 1 ones, although some degree of admixture was still evident between the two populations. Population 1 shows a more diffuse distribution in principal component space, indicating a weaker structure than Population 2, which forms a tighter cluster.

**Figure 6:**
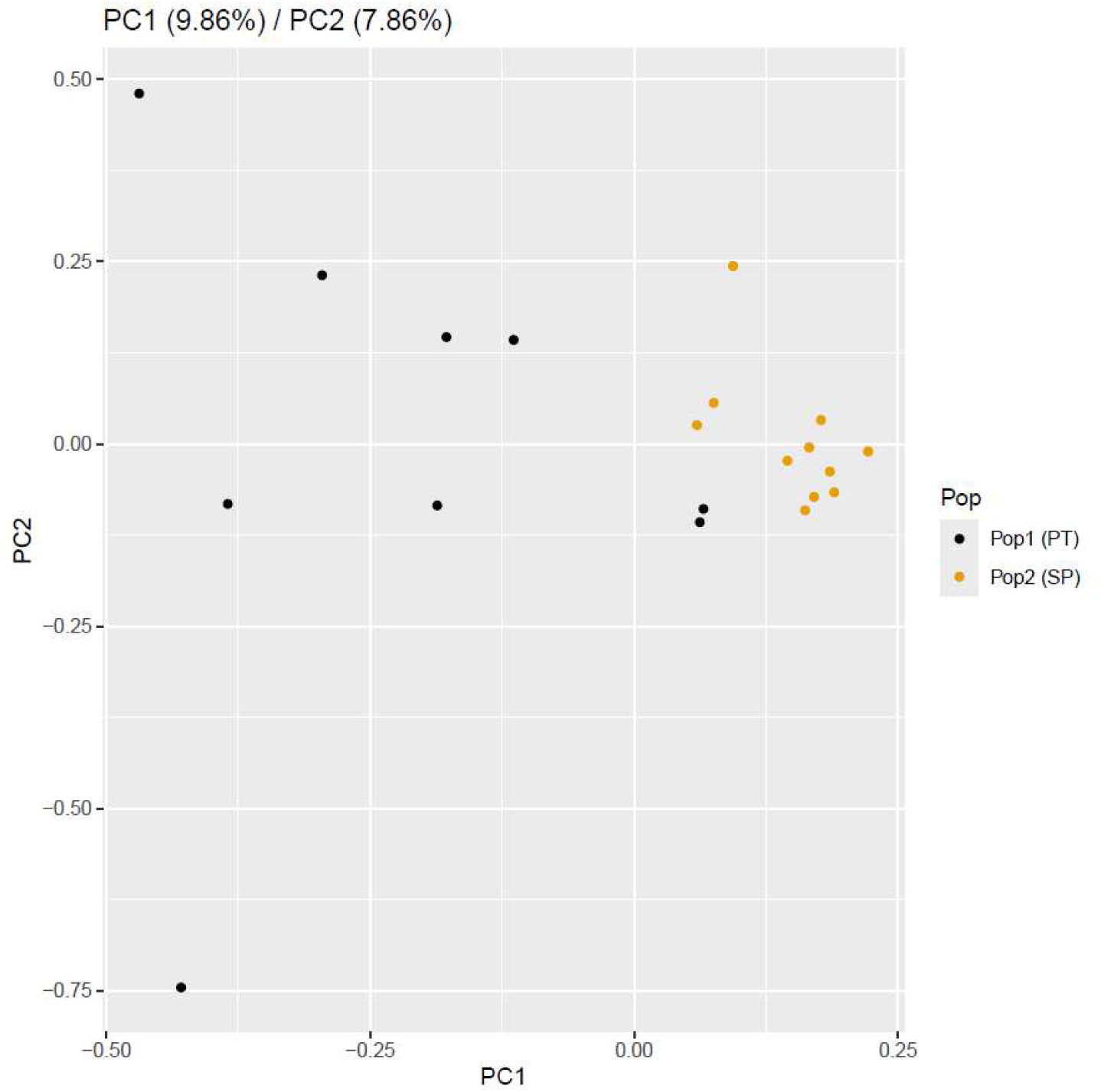
Principal Component Analysis based on genotype likelihood of the genomic regions corresponding to NLRs for the Iberian populations.

## Discussion

### Cork oak immunity is anchored in RNL expansion and NLR diversification

To understand the diversity and function of NLR genes, we developed InterNLR, a custom software tool designed to address the limitations of the existing software (Fernandez-Gutierrez and Gutierrez-Gonzalez, 2021), namely by (i) identifying both canonical and non-canonical NLRs, (ii) recognizing the domain order of NLRs to properly identify each gene, and (iii) detecting RNLs. The maximum likelihood phylogenetic analysis that was conducted with the NB-ARC sequences of both canonical and non-canonical NLRs revealed distinct clades, which emphasizes the high conservation of the NB-ARC domain, allowing to generate clade polarization based on the N-terminal region. The fact that two NLRs were misplaced outside their predicted family can be related to either specific evolutionary pressures or even past recombination events. The RNL clade displayed a well-conserved and polarized pattern compared to the other clades. This observation aligns with the proposal that RNLs primarily function as NLR “helpers”, exhibiting a higher degree of conservation, while “sensor” NLRs are subject to more mutational events driven by selection pressure from new pathogen adaptations (Cesari, 2018). This selective pressure contributes to “birth” and “death” events within the NLR clades, leading to the existence of truncated and non-canonical forms. Notably, within each clade, NLRs with intact N-terminal regions tend to be located adjacently, such as CNs next to CNLs. Comparing with the other two canonical NLR classes, the lower ratio of “number of CNLs” to “CNL-like genes” suggests a potentially more dynamic gene turnover process within this class, resembling a scenario consistent with birth and death evolutionary dynamics (Jacob et al., 2013).

Both the phylogenetic analysis of the three *Quercus* species and the Orthofinder results point towards the occurrence of gene duplications within the RNL clade. This phenomenon may be reason for the higher ratio of “number of RNLs” to “number of TNLs” observed in cork oak compared to the other two species. It suggests that the immune system of cork oak places a greater reliance on effector-targeted host proteins for recognition, often needing the involvement of additional proteins like “helper” RNLs for activation (Saile and Kasmi, 2023). These ratios can vary significantly among different plant species, varying low as 0.04 in *Solanum tuberosum* to high as 0.38 in *Vitis vinifera* (Shao et al., 2016). In tree species, such as seven conifer species, as studied by Ghelder et al. (2019), this ratio can reach 0.10. This range in ratios reflects both the complexity and the species-level specificity strategies that plants employ to combat pathogens.

It’s noteworthy that all RNL genes within the orthogroup displaying gene duplications share orthology with AT5G66900 (N Requirement Gene 1, NRG1) in *Arabidopsis*. NRG1, a subfamily of RNLs closely related to ADR1 in *Arabidopsis*, serves as a “helper”, as demonstrated by Castel et al. (2019). It plays an import role by facilitating the hypersensitive cell death response, ensuring complete resistance against oomycete pathogens in *Arabidopsis*. This study revealed that the involvement of NRG1 in bacterial pathogen resistance in *Arabidopsis* is not essential since its function is shared with ADR1 (Castel et al., 2019). Therefore, examining both the regulation and the expression patterns of NRG1 orthologues in response to biotic stresses will deepen our understanding of their roles in plant defence mechanisms.

### Expression profiles unveil functional and evolutionary layers of NLR diversity

Analyses of NLR gene expression in 1.5-year-old stems revealed distinct patterns, suggesting tissue-specific specialization. Differential analysis between tissues highlighted a greater diversity of DE NLRs with high expression in xylem compared with phellem and inner bark. The high diversity of NLRs expressed in the xylem may reflect the need for redundancy and collaboration among different receptor classes to secure this vital tissue. Since the xylem is essential for water and nutrient transport and is a primary target of vascular pathogens and environmental stresses, multiple NLR classes acting in concert could provide a broader recognition spectrum and a more robust immune response. Additionally, when looking to the expression profiled by NLR class we found that there was significantly higher expression of RNL genes in the xylem compared to other classes, suggesting a potential specialization of this particular tissue in the “sensor”/“helper” system (Saile et al., 2020).

In stems, LOC112003710, an orthologue of RPP13, stood out as the gene with highest expression gene in phellem. In *Arabidopsis*, this gene encodes a CNL. It is known for its ability to confer resistance to various strains of *Peronospora parasitica*, a biotrophic oomycete responsible for downy mildew (Bittner-Eddy et al., 2000). Interestingly, within cork oak, LOC112003710 lacks the typical C-terminal Leucine Rich Repeat (LRR) domain commonly associated with CNLs. However, previous research has suggested that an NLR without the LRR domain can still function as a guard and initiate defence response (Zhao et al., 2015). One gene that the highest expression in inner bark and xylem was LOC112022539, orthologue to *Arabidopsis* ADR1. Recent research suggests that ADR1 (and other RNLs) interact with the plasma membrane (PM) and activate hypersensitive response (Saile et al., 2021).

By analyzing the average percentage of expressed genes for each NLR class and the top five genes with higher mean expression values in each tissue, it becomes evident that the levels of expression of non-canonical NLRs cannot be negligible. While it may be difficult to ascertain specific functions based solely on expression assays, it is reasonable to infer that some of the NLRs may have or have had some function. As said before, these genes may be subjected to high selection pressure that comes from the pathogen adaptation, and this pressure result in a high degree of birth and death of NLRs (Monteiro and Nishimura, 2018). In this context, some non-canonical forms may represent remnants of formerly functional genes whose promoters remain active. Conversely, other forms may represent emerging NLRs in the process of acquiring function. Conducting a promoter profiling study would be crucial to further investigate and address this issue.

Previous studies have highlighted the presence of a basal level of expression for most NLRs, ensuring that the plant is prepared to counteract pathogen invasions that surpass the initial line of defence provided by PAMP-mediated triggered immunity (Jones et al., 2016). However, our investigation uncovers significant differential expression patterns of numerous NLRs under drought conditions in different tissues, including phellem, inner bark, and xylem. Particularly in the xylem, 97 canonical and non-canonical NLRs exhibited differential expression, indicating the critical role of this tissue in plant survival and adaptation to abiotic stresses.

The extensive literature review aiming to identify genes regulated by abiotic stress, together with the orthologue search within cork oak NLR-ome, opened way for a functional analysis of these genes. Out of the total 918 canonical and non-canonical NLRs, 27 were identified as orthologues to genes present in the list of abiotic stress-regulated genes identified in the literature (Supplementary Table 6). Among these, 7 displayed differential expression under drought conditions. From these genes, LOC112031821 (TN) is orthologue of the ACQOS locus, which includes AT5G46490, AT5G46500, AT5G46510, and AT5G46520. This locus has been previously identified as to be up-regulated in response to osmotic stress in root tissues.

However, our study demonstrated different results. We observed down-regulation of this gene in phellem and inner bark under drought stress conditions. It’s worth noting the differences in our assay compared to the study by Ariga et al. (2017), since we focused on stem tissues and induced drought stress, while their study focused on root tissues and osmotic stress. These differences may be sufficient for the variation on gene expression patterns observed. Furthermore, LOC111996925 (N) and LOC112006457 (TNL) are orthologous to AT5G45490 and AT5G36930, respectively. Intriguingly, all these genes, except LOC111996925, displayed down-regulation in inner bark tissues. This pattern aligns with the findings of Yang et al. (2021), who reported that NLR genes, along with other genes induced by biotic stresses, are often repressed by abscisic acid, high temperatures, and drought. These results suggest that the transcriptional regulation of NLR genes may play a pivotal role in mediating the crosstalk between abiotic and biotic stress responses.

Another gene, LOC111996439 (CNL), displayed differential expression under water stress conditions, specifically showing up-regulation in the xylem tissue. This gene is an orthologue to *Picea glauca* PG_003080_T.1 and *Arabidopsis* AT3G14470, encoding a putative disease resistance RPP13-like protein. Moreover, PG_003080_T.1 has been reported to be up-regulated in response to drought, aligning with our findings and underscoring its potential role in abiotic stress response (Ghelder et al., 2019). Additionally, LOC112040104 (NL) emerged as another gene with differential expression patterns under water stress conditions, up-regulated in both inner bark and xylem tissues. This gene is orthologue to *Arachis hypogaea* 0MH239.1 and WIN0ZV.1. Both have been previously reported to be up-regulated in response to drought conditions (Rizwan et al., 2023). Moreover, it is orthologous to Arabidopsis RPM1, a CNL protein that confers resistance to *Pseudomonas syringae* that induces hypersensitive response (Boyes et al., 1998). These findings suggest the potential role for LOC112040104 (NL) in drought stress responses, possibly involving mechanisms similar to those observed in Arabidopsis RPM1-mediated resistance against bacterial pathogens.

Regarding the remaining gene from the seven with differential expression under drought conditions orthologues to abiotic stress-related genes identified in the literature, LOC112022539 (RNL), it shares orthology with *Arachis hypogaea* 3L0H24.1, which has been reported to exhibit up-regulation in drought conditions (Rizwan et al., 2023). Similarly, *Arabidopsis* ADR1, an LOC112022539 orthologue, has been linked to enhanced drought tolerance when overexpressed in the plant (Chini et al., 2004). Our findings align with these observations, as LOC112022539 also shows up-regulation in xylem tissues during drought conditions. This consistency in up-regulation across different plant species, including the orthologue of this gene in *Abies alba*, reinforces the potential significance of LOC112022539 in mediating drought responses at xylem level in cork oak (Behringer et al., 2015), thus playing an important role in edapho-climatic conditions favoring prologued drought episodes.

### One CNL can have double function in cork oak

The analysis of the *Phytophthora cinnamomi* transcriptomic assay highlighted specific NLRs that may respond to this stimulus in root tissues (thin white roots from 2-month old plants and roots from germinating acorns). When analyzing the genes with higher mean expression values in this assay, LOC112022539 (RNL), ADR1 orthologue, was also highly expressed, suggesting that it can also have a basal constitutive expression in roots. Two other genes, LOC112003710 (CN) and LOC112001962 (NL), orthologues to RPP13 and AT1G50180, respectively, were found to display the same pattern, being highly expressed in both root and stem tissues, likely indicating constitutive expression.

The highest ratios of transcript values between infected and non-infected roots may provide cues about which genes could be up-regulated upon infection. In this case, the highest ratio was observed for LOC112017093 (CN), a RPM1 orthologue. In *Arabidopsis*, this gene is required for hypersensitive response to *Pseudomonas syringae* infection (Boyes et al., 1998). Yet, it is difficult to extrapolate an identical mode of action in cork oak response to *P. cinnamomi*, considering the very different biological systems.

LOC111996439 (CNL) and LOC112016116 (RNL) were found induced in roots from germinating acorns infected with *P. cinnamomi*. LOC112016116 shares homology with *Arabidopsis* AT5G66900 (NRG1) a “helper” gene required for hypersensitive cell death response (Castel et al., 2018). This gene is also included in the RNL orthogroup where gene duplications were detected in cork oak. In this context, the gene duplications within this orthogroup in cork oak may indicate a diversification of RNL genes, potentially resulting in functional differences or adaptations to specific environmental or pathogenic pressures. However, to assess if this diversification contributes to functional innovation or merely reflects redundancy, we would need further functional genomic and evolutionary analyses. LOC111996439 (CNL) showed the highest ratio of induction in early developmental stages, meaning that it may be particularly important for plant survival when the germinating acorn is infected with the pathogen. This gene was also more expressed in drought conditions, a behaviour also observed for its orthologue (PG_003080_T.1) in *Picea glauca* (Ghelder et al., 2019). This dual function suggests a central role of LOC111996439 in cork oak, contributing to the plant protection mechanisms in both biotic and abiotic stresses.

### Genetic diversity and evolutionary dynamics of NLRs

The evolution of NLRs is shaped by the constant interplay between plant hosts and pathogens, leading to distinct evolutionary strategies (Stam et al., 2016). Yet, the evolutionary dynamics between pathogen effectors and host NLRs remains a subject of ongoing debate. The co-evolutionary Red Queen ‘Arms Race’ model states that the genomes of both pathogen and plant evolve to overcome defences or avoid infection, respectively, suggesting that positive selection acts on NLRs (Bakker et al., 2006). The alternative model – ‘Trench Warfare’ – predicts that there is long-term accumulation of genetic variation in the plant host that allows it to keep pace with parasite effector variations, suggesting balancing selection acting on NLRs (Stahl et al., 1999, Tellier et al., 2014).

Selection signatures can be elucidated through population genetic summary statistics calculated on NLR regions. The fact that we obtained a lower mean observed heterozygosity (HO) than mean expected under Hardy-Weinberg Equilibrium (HE, assuming a population that is randomly mating, with no selection, migration, or genetic drift) may indicate a deviation from neutrality. Although this pattern is commonly associated with purifying selection or inbreeding, it can also indicate the action of recent positive selection. In such cases, a selective sweep may drive a beneficial allele to high frequency or fixation, reducing genetic diversity and observed heterozygosity in the surrounding genomic region (Watterson, 1977; Payseur et al., 2020). Low mean nucleotide diversity (π), as observed, is also an indicator of positive selection (Noll et al., 2022) – when a beneficial allele rapidly increases in frequency, linked variation in the surrounding region is reduced, leading to a local decrease in genetic diversity.

Both mean observed heterozygosity and nucleotide diversity are relatively low, which may reflect the action of recent positive selection, consistent with the Red Queen ‘Arms Race’ model. However, this interpretation appears to contrast with the positive mean Tajima’s D observed in NLRs.

Tajima’s D evaluates the deviation between nucleotide diversity (π) and the number of segregating sites (Watterson’s θ) (Tajima 1989). When π exceeds θ, Tajima’s D becomes positive, often indicating balancing selection. Conversely, when π is lower than θ, Tajima’s D turns negative, which may suggest either purifying or recent positive selection (Carlson et al., 2003).

It is important to note, however, that the positive mean Tajima’s D obtained may be disproportionately influenced by a small number of positive outlier NLRs with exceptionally high values (see Table 7). These loci could be under strong balancing selection, artificially inflating the mean and masking a broader trend of neutrality or the more directional selection across the majority of genes.

Therefore, while the low diversity and heterozygosity obtained may be consistent with selective sweeps, the overall positive Tajima’s D highlights the possibility that long-term maintenance of allelic diversity is occurring in at least a subset of NLRs. Also, there is also a considerable number of NLRs that, individually, either have positive or negative Tajima’s D values. Taken together, these observations suggest that both evolutionary models – ‘Arms Race’ and ‘Trench Warfare’ – may be acting simultaneously across different NLR loci.

Principal Component Analysis based on NLRs revealed a clear separation of the two Iberian populations, the Spanish (Population 2) and the Portuguese (Population 1), although with signs of admixture between them. This suggests that both genetic differentiation exists, and gene flow and/or shared ancestry may have contributed to the observed pattern. Moreover, the wider spread of the Portuguese population in the PCA space matches its higher nucleotide diversity. This is supported by a larger π obtained for the Portuguese population, suggesting higher pairwise genetic differences among individuals.

### Conclusions

In this study, we employed InterNLR, a novel tool for precise NLR gene annotation, to unravel the complexity of the NLR gene family in *Quercus* sp. We identified 918 NLR genes in cork oak, encoding canonical NLRs (CNLs, TNLs, and RNLs) and non-canonical NLR-like proteins. Phylogenetic analysis, particularly focused on the NB-ARC domain, unveiled the evolutionary relationships among NLR clades. Our findings support the ancestral nature of TNLs and the specialized role of RNLs as NLR “helpers”. Gene duplications were detected in the latter clade.

We observed distinct expression patterns of NLRs across various cork oak tissues, supporting tissue-specific functional roles. In particular, RNLs displayed significantly higher expression levels in xylem, suggesting a specialization in this vital tissue.

Seven NLRs exhibited differential expression under drought conditions and shared orthology with genes known to respond to abiotic stress. One gene in specific, LOC111996439, a CNL, was part of this seven NLRs and is possibly up-regulated on young roots infected with *Phytophthora cinnamomi* zoospores. This dual functionality identification in responding to both biotic and abiotic stresses is indicative of its central role in early defence mechanisms. The identification of LOC112022539, ADR1 orthologue, as highly expressed in both transcriptomic assays suggests a baseline expression and a potential crosstalk between abiotic and biotic stress responses mediated by this gene. Lastly, the differential expression of LOC112016116,

NRG1 orthologue and its presence within the RNL where gene duplications have been detected raises intriguing questions about the functional significance of these duplications and the potential role of NRG1-like genes in cork oak’s immune responses. To strengthen our conclusions, further functional studies complementing *Arabidopsis* mutants with cork oak genes of interest could provide further insights into their functional roles.

Furthermore, the detection of both balancing and positive selection across distinct NLRs supports the hypothesis that the evolution of the NLR-ome may be driven by a combination of Red Queen ‘Arms Race’ and ‘Trench Warfare’ models, providing a reconciliatory perspective on the long-standing debate regarding the evolutionary dynamics of host-pathogen interactions. Along with the clear population structure detected between Portuguese and Spanish populations, the diversity revealed in the NLR repertoire not only advances our understanding of cork oak immunity and evolution but also provides a valuable genomic resource for conservation and breeding strategies in forest trees.

## Materials and Methods

### Identification of *Quercus* NLRs

InterNLR is a software tool developed in this study, to overcome the limitations of available tools in the identification of (non-/)canonical NLRs, recognition of domain order in NLR for proper gene identification, and detection of RNLs. Its development and running used predicted protein sequences from *Quercus suber* (assembly GCA_002906115.5, Usié et al., 2023), *Quercus lobata* (GCA_001633185.5, Sork et al., 2016) and *Quercus robur* (GCA_932294415.1, Plomion et al., 2018) as inputs. This Bash wrapper firstly uses Interproscan (Zdobnov and Apweiler, 2001) analysis against ‘gene3d’, ‘pfam’ and ‘COILS’ databases. After this, NLR domains are called by *domain_caller.py*, which creates a list of all domain matches and their respective gene coordinates. This list is then lastly used as input for *nlr_caller.py* to obtain the final output, consisting of several plain text files containing the gene IDs according to each NLR type. *Q. suber* and Q. *lobata* files included all alternative splicing variants, so for each genomic position (LOC id) a splicing variant (XP id) was arbitrarily chosen as representative. InterNLR is available on GitHub – https://github.com/lmgoncalves94/internlr.

### Phylogenetic analysis of NB-ARC domains

The coordinates of the NB-ARC domain for each NLR were retrieved according to ‘gene3d’ and ‘pfam’ databases (from Interproscan) and Bedtools 2.18 (Quinlan and Hall, 2010) was used to obtain the peptide sequences corresponding to the NB-ARC domain of each NLR. NB-ARC sequences were aligned using the MAFFT Version 7 GUI with default parameters (https://mafft.cbrc.jp/alignment/server/, Katoh and Toh, 2008). The resulting alignment file was trimmed in TrimAl 1.3 (Capella-Gutiérrez et al., 2009) with a gap threshold of 0.9 and a minimum percentage of the positions in the original alignment to conserve of 0.6. The trimmed alignments were used as input for RAxML 8.0.0 (Stamatakis, 2014) to construct a maximum-likelihood phylogenetic tree, with bootstrap analysis (1000 replications). The consensus trees were edited and plotted using FigTree 1.3.0 (http://tree.bio.ed.ac.uk/software/figtree/). Orthofinder was run to detect gene duplications (Emms and Kelly, 2019).

### Gene expression analysis

The gene expression analysis started with an already existing dataset of RNA-seq raw read counts of three tissues of cork oak. The experimental design consisted on one-year old plants that were maintained in two contrasting irrigation conditions for 6 months (March to August). Tissue-specific sampling was conducted by collecting 10 cm segments above the root collar, specifically targeting the xylem, inner bark, and phellem tissues. Raw read counts were retrieved from ArrayExpress (accession E-MTAB-13376). TPM values were calculated from raw read counts in the R environment, and genes with TPM ≥ 1 were considered expressed. Differential expression analysis between the three tissues (independently of growth conditions) and between WW and WD was conducted using DESeq2, as described in Barros et al. (2024).

The publicly available transcriptomic dataset consisting of RNA-seq data of roots on different developmental stages - roots from germinated acorns (days old) and thin white roots (2 months old) - both infected and non-infected with *Phythophthora cinnamomi* zoospores was retrieved from CorkOakDB.

### Variants call

Raw RNA-seq sequencing reads were downloaded and converted to FASTQ files using sra-toolkit (Leinonen et al., 2010) for each SRA from each of the two bioprojects – PRJEB33874 and PRJNA650215. Trimmomatic (Bolger et al., 2014) was used to remove adapters and low-quality (Q < 15) and small (read length < 36 bp) reads. STAR (Dobin et al., 2013) was run to map the sequencing reads to the reference genome and obtain BAM files. SAMtools (Danecek et al., 2021) was used to remove low-quality mapped reads (q < 20). Mapped reads were arbitrary removed to normalize the coverage for each BAM file (mean target coverage around 20 X). After normalization, FeatureCounts (Liao et al., 2014) was ran to obtain the raw read counts per site.

Only NLRs (canonical and non-canonical) with an average of at least 10 mean raw counts per gene were selected for future analysis, corresponding to constitutively expressed NLRs. Those genes were filtered with BEDtools and BAM files with only those NLR regions were obtained. After filtering, Samtools was run to sort and index each resulting BAM file.

To call variants within NLR regions from RNA-seq, BCFtools (Li, 2011) was employed. Given that the analysis focused on constitutively expressed regions, we treated these as representative of the genomic sequence. Variants were filtered using quality thresholds (QUAL ≥ 20) and depth (DP ≥ 10) to ensure reliability.

### Population genetics and structure

To obtain the population genetics statistics, VCFtools (Danecek et al., 2011) was used. It was run for all constitutive NLR regions with --het option to obtain the observed and expected homozygosity, which were used to obtain the weighted mean observed (H_O_) and expected heterozygosity (H_E_) under Hardy-Weinberg Equilibrium using an in-house custom python script. Nucleotide diversity (π) and Tajima’s D were calculated in 1000 bp sliding windows with --window-pi 1000 and --TajimaD 1000 options, respectively. Both weighted mean π and Tajima’s D were obtained for all NLRs using custom python scripts. Another in-house script was also run to obtain the Tajima’s D per NLR.

ANGSD (Korneliussen et al., 2014) was used to compute genotype likelihoods and allele frequencies for PCA analysis. The tool was run with the parameters: -doCounts 1, -GL 1, - doMajorMinor 1, -doMaf 1, -skipTriallelic 1, -doGeno 32, and -doPost 1. ngsCovar (part of NGS Tools; Fumagalli et al., 2014) was used to compute the covariance matrix for principal component analysis (PCA) based on genotype posterior probabilities. The input genotype likelihood file (.geno), generated by ANGSD, included 20 individuals. Parameters included -call 0 to use genotype posterior probabilities directly, and -norm 0 to avoid normalization of the covariance matrix. The covariance matrix produced by ngsCovar was used as input to generate the principal component plots using the plotPCA.R script from the NGS Tools package (Fumagalli et al., 2014). The first two principal components (PC1 and PC2) were plotted, and individuals were grouped according to a population metadata file (.clst).

## Supplementary Data

**Supplementary Figure 1: Maximum Likelihood Phylogeny of *Q. suber, Q. lobata* and *Q. robur* canonical NLRs based on the NB-ARC domain.** Colors indicate NLR sub-groups: Red – TNLs; Green – CNLs; Blue – RNLs. The NB-ARC domain of *Physcomitrium patens* PRG gene was used as outgroup (purple). It is possible to distinct the polarization of each of CNL, TNL and RNL clades. The phylogenetic tree scale represents substitutions per site. Branch support values result from bootstrap analysis (1000 replications).

**Supplementary Figure 2: Maximum Likelihood Phylogeny using the NB-ARC domains of the 918 NLRs and overview of gene expression patterns in different layers of cork oak stem.** a – NLR type per phylogenetic position (NL – black, TNL – red, CNL – green, RNL – blue, TCNL – yellow, RTNL – magenta, N – grey, TN – light red, CN – light green, RN – light blue, *Physcomitrium patens* PRG – purple). b – Classes of NLRs (RNL-like – blue, TNL-like – red, CNL-like – green). c – NLRs of interest, namely orthologues to abiotic stress-induced NLRs (grey, genes related to abiotic stress response in other plant species), genes both orthologues to abiotic stress-induced NLRs and differently expressed under drought stress (red). d – Differential expression pattern (log_2_ fold change) of NLRs under water-deficit conditions, from blue (down-regulated) to red (up-regulated) in: d1 – phellem; d2 – inner bark; d3 – xylem. e – Overall mean expression (TPM) in the three stem tissues, under well-watered and water deficit conditions: phellem (e1), well-watered - e1a, water-deficit - e1b; inner bark (e2), well-watered - e2a, water-deficit - e2b; and xylem (e3), well-watered - e3a, water-deficit - e3b. The phylogenetic tree scale represents substitutions per site. Branch support values result from bootstrap analysis (1000 replications).

**Supplementary Table 1:** Orthogroups identified through Orthofinder analysis for TNLs.

**Supplementary Table 2:** Orthogroups identified through Orthofinder analysis for CNLs.

**Supplementary Table 3:** Orthogroups identified through Orthofinder analysis for RNLs.

**Supplementary Table 4:** NLRs exhibiting duplications as identified through Orthofinder analysis, supplemented by blastp comparisons against the *Arabidopsis* proteome.

**Supplementary Table 5:** Number of expressed genes and its percentage per tissue in cork oak stems.

**Supplementary Table 6:** NLRs identified as related to abiotic stress response across several plant species with orthologues in cork oak.

